# The Impact of New Urbanization on Quantity Increase and Quality Improvement of Green Innovation: Based on Multivariate Moderating Effect Models

**DOI:** 10.1101/2025.07.13.664607

**Authors:** Liang Fang, Chengxiang Li, Chen Jin

## Abstract

New urbanization is a major national strategy that China has proposed and implemented. Whether new urbanization (NEU) can promote quantity increase and quality improvement of green innovation (GI) is a common concern of the government and scholars. This paper empirically analyzes quantity increase and quality improvement effects of new urbanization on urban GI by adopting the panel data of each city from 2011 to 2022. The study finds that new urbanization not only has a positive contribution to the quantity increase of GI in cities but also plays a positive role in the quality improvement of GI. Digital inclusive finance will further amplify the dividends of new urbanization and exert the GI effect. The interaction of digital inclusive finance and government support, informatization, regional economic development, and energy consumption also amplify the dividends of NEU and exert strong “quantity increase” and “quality improvement” effects on GI. The stronger the government’s green support, the more conducive it is to new urbanization to further exert quantity increase and quality improvement effects on GI. The conclusions of this paper have important policy implications for further promoting the construction of new urbanization and GI.

## 1. Introduction

Over the past decade, China’s urbanization has progressed swiftly, resulting in a significant influx of rural residents into metropolitan areas, which has catalyzed extraordinary urban development and the emergence of multiple city clusters. By virtue of their strong industrial agglomeration effect and innovation and R&D capability, city clusters have become economic growth poles [1]. Traditional urbanization has produced a series of drawbacks, with over-consumption of resources and environmental pollution being prominent [2], and some cities are overly reliant on traditional low-end industries or labor-intensive industries, leading to a lack of demand for scientific and technological innovation. China has put forward a major strategy of new urbanization (NEU), which no longer pursues the expansion of city scale and increase in population but emphasizes the upgrading of urban functions, the elevation of public service standards, and the achievement of green and low-carbon transformation [3]. “The State Council Government Work Report 2025” proposes to promote NEU in a scientific and orderly manner, adhere to innovation-led development, promote technological innovation, and build up basic and strategic support for Chinese-style modernization. China is actively exploring how to optimize the structure and allocation of resources and facilitate the low-carbon transformation of industries by advancing NEU so as to embark on the road of green and sustainable development. Simultaneously, global competition in the green industry is fierce. The green energy industry is undergoing structural changes, and it is imminent to promote steady economic recovery and healthy development. Under the double pressure of developing economy and ecological environment, it is of great significance to promote the green and low-carbon transformation of the economy and society through NEU and green innovation (GI) and to improve the level of green technological innovation in order to realize sustainable development. In this regard, it is important to explore whether NEU can further inject new kinetic energy into urban green technological innovation so as to realize the win-win situation of economic, social and ecological benefits, which will become an important breakthrough in solving urban development problems.

Existing literature research shows that NEU not only effectively diminishes pollutant emissions and enhances energy efficiency but also exerts a substantial influence on ecological effect [4]. NEU realizes the concentration of population and resources in urban regions, which facilitates the enhancement of industrial structure, regional integration, and economies of scale, thereby advancing urban economic growth. [5]. Some scholars focus on the linkage between urbanization and ICT and the study of digital divide issues. Wang Zhaohua et al. (2019) suggested that NEU indirectly mitigates CO2 emissions by eradicating the rebound effect of energy-efficient technologies on CO2 emissions and fostering environmental technologies, resulting in a threshold effect on CO2 emissions based on the disparity between the levels of energy-saving and environmental technologies. [6]. Literature has also focused on the impact of urbanization patterns on the stages of the innovation process and regional spillovers. Tripathi Sabyasachi et al. (2022) propose that urbanization positively influences the phases of the regional innovation process (knowledge creation, knowledge implementation, and innovation production), with the effects of various urbanization patterns differing by stage [7]. Zhang Ran et al. (2023) suggested that the implementation of NEU significantly promotes green technological innovation by alleviating the financial constraints of enterprises, thus positively affecting green environmental protection [8]. Focusing on urbanization innovation policy research to address regional development issues Focusing on urbanization policy and the enhancement of innovation capacity, the literature explores its heterogeneity and mechanism of action. Li Shengsheng et al. (2023) suggest that the pilot policy of NEU significantly enhances urban innovation capacity by enhancing the level of human capital as a theoretical mechanism, which makes the impact of NEU policy on urban innovation at both the city level and regional level both show obvious heterogeneity, especially in big cities and the western region, the impact effect is more significant [9]. Chen Jian et al. (2020) suggested that although the urbanization process enhances the innovation capacity of the region, it exhibits a negative correlation with the innovative capacity of the adjacent regions [10]. Ma Cankun et al. (2022) and Razzaq Asif et al. (2022) focused on GI and NEU research to address synergistic development. The focus is on the synergistic effects of GI and NEU development, exploring its impact on environmental quality and resource consumption [11,12].

The contributions of this paper are: (1) This paper focuses on the relationship between NEU and GI, empirically analyzes the impact of NEU on GI, and specifically discusses the “quantity increase” and “quality improvement” effect of NEU on GI, which enriches the research results of NEU. (2) This paper confirms the impact of digitalization on urban GI, confirms that digital inclusive finance affects the development effectiveness of NEU, points out the development direction of NEU, and provides important policy insights for further promoting NEU. (3) This paper constructs a multivariate moderating effect model. It confirms that the interaction between digital inclusive finance and governmental, technological support, informatization, regional economic development, and energy consumption affects the innovation effect of NEU. The multivariate moderating effect helps to reveal more comprehensively and in-depth the inherent logic and complex mechanism of NEU and GI, which provides a new way of thinking and direction for the subsequent research. (4) From the standpoint of government green support, this paper verifies that the relationship between NEU and GI is not a linear trend, which further enhances comprehension of the complex influence relationship between NEU and GI.

## 2. Research hypotheses

“Urbanization” denotes the transformation of rural populations into urban ones while also encompassing the necessity for robust industrial and social growth. This necessitates the incorporation of ecological civilization concepts and principles throughout the urbanization process, pursuing an intensive, intelligent, green, and low-carbon New Urbanization (NEU) fundamentally guided by a scientific perspective on development.

Firstly, from the perspective of resource agglomeration, NEU emphasizes the transformation of human-centered citizenship [13], which is conducive to the acceptance of higher education and skills training for the transferring population [14] and can cultivate a large number of technicians in the fields of new energy and intelligent manufacturing, create the conditions for “quantity increase” of innovation in human resources. NEU also promotes the construction and industrial layout of urban areas, attracting more capital and resources to concentrate in urban areas [15]. Under the requirements of greening and high-quality development, venture capital, bank loans, and government support funds are shifted to GI projects, which assist firms in increasing R&D investment and carrying out more GI activities. From the perspective of market demand, NEU drives the increase in income level and the change of consumer attitudes [16]; people will increase the demand for green products, energy-saving products and high-tech products, thus triggering the derived demand for GI. Enterprises will increase investment in GI and develop more green products to meet market demand. NEU promotes the development of urban construction, transportation, energy supply and other related industries [17], which will expand the demand for green technologies and solutions. Enterprises will actively develop green building technologies and increase the quantity of GI.

Secondly, urbanization promotes the spatial transfer of resources and factors [18]. At the same time, NEU “abandons” the “quantity” change of traditional factor transfer and puts forward higher requirements for the quality of resource and factor transfer. NEU will be more inclined to attract high-level and high-skilled talents to urban agglomeration [19]. High-quality human capital promotes technological research and development through “learning by doing” [20], realizing the transformation of innovation from low-end imitation to original breakthroughs, which promotes the improvement of the quality of GI. NEU promotes the integration and development of different industries and fields [21], which brings more technological cross-fertilization and integration opportunities for GI. Information technology, energy technology, and environmental technology are integrated, which improves the technical content and practicality of GI achievements. NEU policy puts forward the requirements of GI and formulates strict environmental protection standards and industry norms, which promote enterprises to improve the quality of GI.

### Therefore, hypotheses 1: NEU is conducive to promoting the “quantity increase” and “quality improvement” of GI

The development of digital inclusive finance is crucial for improving the accessibility of financial services to the actual economy, facilitating the equitable distribution of financial resources, and fostering high-quality economic development. Relying on technologies such as big data, blockchain, and artificial intelligence, digital inclusive finance breaks through the dependence of traditional finance on collateral, offering accessible financing avenues for small-sized enterprises, individual entrepreneurs, and other urban entities [22]. However, in reality, the level of digital inclusive finance development in different regions varies greatly, and cities exhibiting advanced economic development generally possess enhanced digital inclusive finance progress because the advancement of digital inclusive finance is intricately linked to both economic development and information technology development [23]. Some scholars have found that in areas with better digital inclusive finance, it is easier for migrant populations to obtain financial support for rent, education and living and for laborers to obtain funds for employment and entrepreneurship [24], which help them integrate into urban life. Digital inclusive finance is also believed to channel funding towards green industries, emerging fields of smart manufacturing, ecological restoration, and clean energy projects, which are injecting significant impetus for the advancement of green industries [25]. Moreover, digital inclusive finance improves the efficiency of matching green technology and market demand through the information-sharing mechanism. It promotes collaborative innovation within the upstream and downstream sectors of the green industry chain [26]. Conversely, suppose the advancement of digital inclusive finance in the region is deficient. In that case, it affects urban construction, industrial upgrading, and population movement in urbanization [27], especially the lack of digital financing tools for green industries and eco-city construction projects, which makes it difficult to attract social capital. It is also difficult for clean energy and ecological restoration projects to obtain targeted financing.

### Therefore, hypothesis 2: the higher the level of digital inclusive finance, the more conducive to the “quantity increase” and “quality improvement” of NEU on GI

In addition to the impact of digital inclusive finance, government technology support also has an important impact on GI and NEU. The government is an important promoter of urban construction and development, and it can not only formulate policies and regulations but also provide support for urbanization and S&T innovation through appropriations, establishment of funds, tax incentives, and risk compensation [28]. As the complexity of technological innovation increases, the government’s technological support will play an increasingly significant role in stimulating the main role of enterprises and enhancing the effectiveness of urban governance [29].

The swift advancement of the global economy has rendered informationization a crucial catalyst for societal progress. Information technology can deeply intervene in the development of green energy materials, expedite the research and development cycle, diminish research and development expenditures [30], and accelerate the development and iteration of green technology. With the help of information technology, enterprises can realize the intelligence of the production process, reduce the errors and waste of resources caused by manual operation, and improve the efficiency of R&D [31]. The level of regional economic development also affects NEU and GI. Economically advanced regions typically possess greater financial resources for the research, development, and implementation of green technology infrastructure. They can increase investment in renewable energy technology and environmental conservation sectors to promote the output of GI results [32]. Regions with good economic development are more attractive to high-quality talents, providing intellectual support for GI. There are substantial disparities in energy use among various cities. In general, high-energy-consuming cities tend to face greater pressure on energy consumption and environmental pollution [33], and this pressures cities to prioritize GI in the process of NEU. In order to reduce energy consumption and pollution, these cities will actively promote R&D and implementation of sustainable technologies, thus creating a strong demand for GI. NEU underscores the principles of sustainable development and green ecology, advocating for the transition of urban energy systems from high-carbon to low-carbon alternatives. Cities with high energy consumption will prioritize the production and exploitation of ecological energy in the NEU process, thereby reducing reliance on traditional energy sources, decreasing carbon emissions[34], and promoting the innovation and development of green energy technologies.

### Therefore, Hypothesis 3: the higher the level of governmental and technological support, informatization, regional economic development, and energy consumption, the stronger the moderating effect of digital inclusive finance on NEU and GI

## 3. Research design

### 3.1. Indicator design

#### i) Dependent variable

The dependent variable is GI. To reflect the “quantity increase” and “quality improvement” of GI, this study uses the total number of green patent applications (GIN) per 10,000 people to measure the “quantity increase “of GI and the number of green invention patent applications (GNS) per 10,000 people to measure the “quality improvement” of GI [35].

#### ii) Independent variable

The independent variable is NEU. According to its connotation and characteristics, NEU includes population urbanization, economic urbanization, environmental urbanization, and social urbanization [36]. The indicators are shown in Table 1, in which the ratio of urban-rural Engel’s coefficient, the ratio of urban-rural consumption, and the ratio of urban-rural income are negative metrics and are taken as reciprocal. The multivariate indicator system uses the entropy weighting method to reduce the dimensionality processing to get the comprehensive score.

**Table 1.** Indicators of NEU.

| First-level Indicators | Second-level Indicators | Third-level Indicators | Weight |
| --- | --- | --- | --- |
| NEU | Population urbanization | Urbanization rate of permanent population | 0.0203 |
|  |  | Urbanization rate of registered residence | 0.0424 |
|  |  | Urban pension insurance coverage rate | 0.0288 |
|  |  | Number of participants in urban basic medical insurance | 0.1743 |
|  |  | Proportion of general public budget expenditure on social security and employment | 0.0167 |
|  | Economic Urbanization | Population density in urban areas | 0.1663 |
|  |  | The logarithm of economic agglomeration degree | 0.0154 |
|  |  | Per-capita Disposable Income of Urban Residents | 0.0276 |
|  |  | The ratio of Engel coefficient between urban and rural areas | 0.0119 |
|  |  | Urban rural consumption ratio | 0.0090 |
|  |  | Urban rural income ratio | 0.0177 |
|  | Environmental Urbanization | Urban per capita road area | 0.0262 |
|  |  | Urban per capita construction land area | 0.0356 |
|  |  | Per capita built-up area of urban areas | 0.0370 |
|  |  | Green coverage rate of built-up areas | 0.0024 |
|  |  | Harmless treatment rate of urban household waste | 0.0034 |
|  |  | Urban sewage treatment rate | 0.0029 |
|  | Social urbanization | Proportion of hospital hygiene beds to urban permanent population | 0.0177 |
|  |  | Urban water supply penetration rate | 0.0021 |
|  |  | Urban gas penetration rate | 0.0027 |
|  |  | The total collection of books in public libraries | 0.2076 |
|  |  | Proportion of regular undergraduate and junior college students to registered population | 0.1323 |

#### iii) Control variables (CV)

Control variables include the proportion of foreign direct investment to GDP (FDI), science and technology expenditures (TEL), the proportion of total import and export volume to GDP (OOW), the proportion of regular undergraduate and junior college students in the total population at the end of the year (HCL), the Internet penetration rate (INT), the per capita volume of road freight transportation (RFV), and the level of social consumption (SCL).

#### iv) Moderating variables

The moderating variables include the main moderating variable and the grouping moderating variable, the main moderating variable being Digital Inclusive Finance (DIF), data coming from the financial database released by Peking University. The grouping moderating variables are government support for technology (GTS), informatization (INT), per capita GDP (AGDP), and energy consumption (ENC), where GTS is measured by the ratio of science expenditures in the general public budget, the Internet penetration rate measures INT and ENC is measured by the proportion of primary and secondary multiplied by the amount of coal consumed.

### 3.2. Models design

To explore the impact of NEU on GI, a benchmark regression model is designed as:

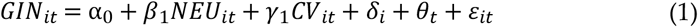

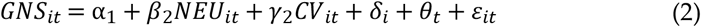

To test the endogeneity issue in the regression model, the two-stage instrumental variable method is employed to construct the 2SLS regression model; NEU lagged one period (LAG_NEU), and nighttime lighting value (NLV) are instrumental variables:

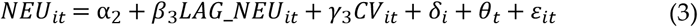

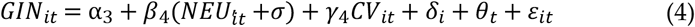

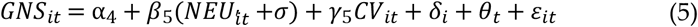

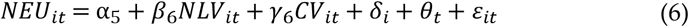

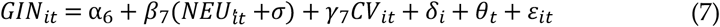

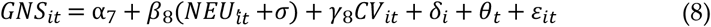

To ascertain the influence of NEU on GI, the interaction term of DIF and NEU (cDIF_it_ × cNEU_it_) is added, and the moderating effect model is constructed as:

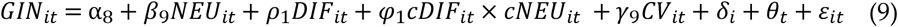

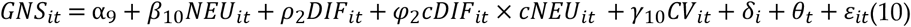

*α* is constant, βis the coefficient of the independent variable, δ is city fixed effect, θ is year fixed effect, and ε is random error.

The data are obtained from the China Urban Statistical Yearbook, statistical yearbooks of various regions, the EPS database, the China Energy Statistical Yearbook, and the China Science and Technology Statistical Yearbook. A few missing values are the result of linear interpolation. The results of the descriptive statistical analysis are shown in Table 2.

**Table 2.** Descriptive Statistical Results.

| VAR | N | Mean | SD | Min | Max |
| --- | --- | --- | --- | --- | --- |
| GIN | 3408 | 0.9645 | 1.5031 | 0 | 13.5925 |
| GNS | 3408 | 0.4246 | 0.7879 | 0 | 9.8401 |
| NEU | 3408 | 0.1209 | 0.0562 | 0.0354 | 0.5371 |
| FDI | 3396 | 0.0148 | 0.0185 | -0.2186 | 0.2827 |
| TEL | 3408 | 0.0174 | 0.0179 | 0.0005 | 0.2068 |
| INT | 3408 | 25.6078 | 14.9261 | 0.3796 | 125.8681 |
| RFV | 3408 | 32.2915 | 65.9670 | 1.2166 | 3042.3968 |
| HCL | 3408 | 0.0200 | 0.0250 | -0.0006 | 0.1502 |
| OOW | 3408 | 0.1758 | 0.2811 | -0.0201 | 2.4913 |
| SCL | 3408 | 0.3817 | 0.1094 | 0 | 1.0126 |
| CIT | 3408 | 0.2312 | 0.0925 | 0.0851 | 0.7392 |
| NGIN | 3408 | 0.6549 | 1.0162 | 0 | 9.9852 |
| NGNS | 3408 | 0.1115 | 0.2471 | 0 | 4.3563 |
| NLV | 3408 | 8.8598 | 9.9050 | 0.1297 | 60.0526 |
| GTS | 3408 | 0.0172 | 0.0173 | 0.0001 | 0.1423 |
| DIF | 3408 | 226.3505 | 83.2703 | 2.7000 | 581.2300 |
| AGDP | 3408 | 5.6743 | 3.3874 | 0.2134 | 25.8482 |
| ENC | 3408 | 6.4742 | 0.7373 | 4.0950 | 8.8560 |
| GES | 3408 | 0.0075 | 0.0039 | 0.0004 | 0.0256 |

## 4 Empirical analysis

### 4.1 Benchmark regression

Whether NEU plays a positive role in the “quantity increase” and “quality improvement” of GI, table 3 shows the results of the benchmark regressions of NEU on GI. E1 and E2 show that the coefficients of NEU on GIN and GNS are 26.9963 and 13.7455, respectively, without controlling the control variables and with controlling year and city fixed effects, which pass the significance test. E3 and E4 show that the coefficients of NEU on GIN and GNS are 26.9963 and 13.7455, respectively; after controlling FDI, TEL, OOW, HCL, INT, RFV, SCL, year fixed effects and cities fixed effects, the coefficients of NEU on GIN and GNS are 23.0471 and 11.9406, respectively, and both pass the significance test. This suggests that NEU not only has a positive effect on the “quantity increase” of GI but also significantly contributes to the “quality improvement” of GI. The comparison of E3 and E4 shows that the “quantity increase “effect of NEU on GI is stronger than the “quality improvement “effect of NEU on GI, and NEU in China is in the development stage of “quantity is stronger than quality” and “transformation from quantity to quality”. NEU has solved the conditions of “consumer demand” and “talent supply” for quantity increase and quality improvement of GI [37]. In addition, NEU advocates the green development mode and high-efficiency development mode, and the strict environmental standards force enterprises to improve efficiency [38] and promote quantity increase and quality improvement of GI. NEU is playing an increasingly significant role in GI through different ways, such as talent pooling, consumption upgrading, and development mode.

**Table 3.** Benchmark Regression Results.

| VAR | E1 | E2 | E3 | E4 |
| --- | --- | --- | --- | --- |
|  | GIN | GNS | GIN | GNS |
| NEU | 26.9963***<br>(5.6124) | 13.7455***<br>(4.7472) | 23.0471***<br>(6.0325) | 11.9406***<br>(5.2395) |
| CV | N | N | Y | Y |
| cons | -2.4344***<br>(-4.8310) | -1.2630***<br>(-4.1675) | -1.2346***<br>(-3.0831) | -0.6372***<br>(-2.7494) |
| *N* | 3408 | 3408 | 3396 | 3396 |
| adj. *R*<sup>2</sup> | 0.434 | 0.307 | 0.519 | 0.410 |
| id FE | Y | Y | Y | Y |
| Year FE | Y | Y | Y | Y |

Note: t statistics in parentheses, \* $p < 0.1$ , \*\* $p < 0.05$ , \*\*\* $p < 0.01$

### 4.2 Robustness test

#### i) Removing special city samples

There are four municipal cities in China: Beijing, Chongqing, Tianjin and Shanghai. In the process of urbanization, these four cities have become the cities that absorb the largest number of rural migrants, which has an important influence on NEU. At the same time, these four cities are also China’s innovation cluster, and the number of innovation achievements is the highest in China. Therefore, the four samples may have an important influence on the innovation effect of NEU. To test the robustness of the regression, this paper removes the samples of the four cities to do the robustness test (table 4). E5 shows that the coefficient of NEU is 23.4855, and E6 shows that the coefficient of NEU is 11.4504, and both meet significance. It shows that the “quantity increase” and “qualitative improvement” effects of NEU on GI are still significant after removing the sample of municipalities, and the regression results are resilient.

**Table 4.** Results of Robustness Test I.

| VAR | E5 | E6 | E7 | E8 |
| --- | --- | --- | --- | --- |
|  | GIN | GNS | GIN | GNS |
| NEU | 23.4855***<br>(6.3638) | 11.4504***<br>(5.2676) | 24.1694***<br>(5.7860) | 12.4691***<br>(5.0389) |
| CV | Y | Y | Y | Y |
| cons | -1.3624***<br>(-3.4681) | -0.6661***<br>(-2.7829) | -1.3445***<br>(-3.1356) | -0.6903***<br>(-2.8038) |
| *N* | 3348 | 3348 | 2830 | 2830 |
| adj. *R*<sup>2</sup> | 0.508 | 0.402 | 0.540 | 0.439 |
| id FE | Y | Y | Y | Y |
| Year FE | Y | Y | Y | Y |

Note: the same as Table 2.

#### ii) Removing special year samples

NEU was proposed as an important strategy in 2015 and was further fully promoted across the country in 2016; in order to prevent these two annual samples from having a disturbing effect on the regression results, this study removes the samples of 2015 and 2016 to test the robustness. E7 shows the regression coefficient of NEU is 24.1694, and E8 shows the regression coefficient of NEU is 12.4691; both pass the significance test. It indicates that after removing the samples from 2015 and 2016, the “quantity increase” and “quality improvement” effects of NEU on GI is still significant, and the regression results are resilient.

#### iii) Adding dummy variables

In addition to NEU, other variables may also have a positive impact on GI. In order to exclude the influence of other variables on the analysis results, this paper controls the role of new-energy model cities policy (NEM) on GI. In this paper, NEM is included as a dummy variable in the regression model for analysis. Cities implementing NEM are 1, and cities not implementing NEM are 0. The test results are presented in Table 5. E9 shows that the coefficient of NEU on the “quantity increase” of GI after adding the dummy variable of NEM is 22.8580, and E10 shows that the coefficient of NEU on the “quality improvement” of GI after adding the dummy variable of NEM is 11.8124, both of which pass the significance test. E10 shows that after adding the dummy variable of NEM, the coefficient of NEU on “quantity increase” of GI is 22.8580, and E10 shows that after adding the dummy variable of NEM, the coefficient of NEU on “quality improvement” of GI is 11.8124, which pass the significance test, indicating that after controlling NEM, the “quantity increase” and “quality improvement” effect of NEU on GI are still obvious. The regression results are robust.

**Table 5.**
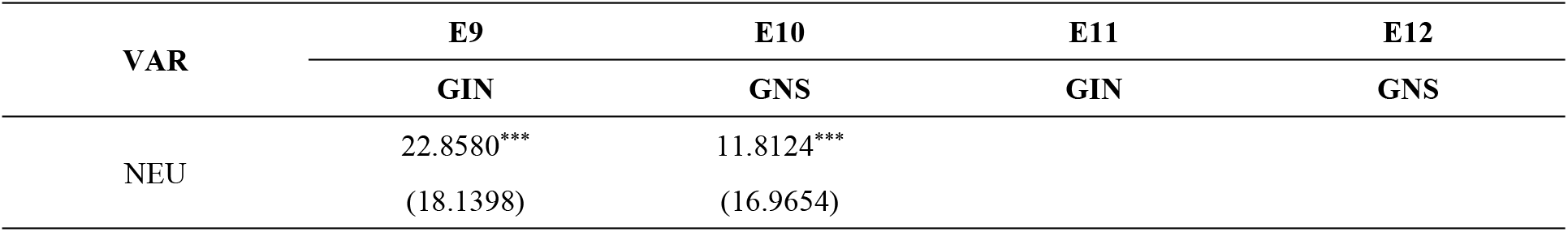
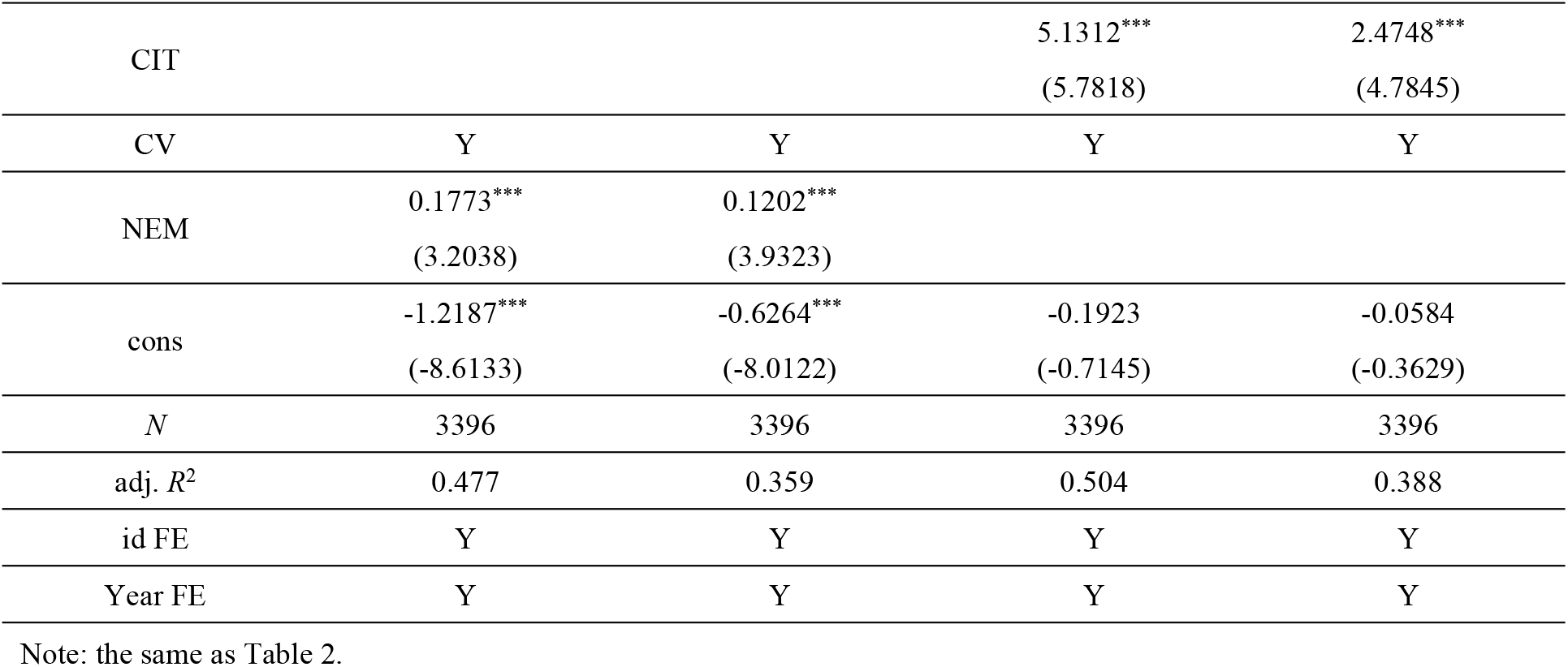
Results of Robustness Test II.

#### iv) Replacing independent variable

Independent variable NEU is a multi-indicator variable; if the indicators of the variable are wrongly selected, it will result in biased regression results. In this study, the citizenship index of the rural transferring population is taken to replace NEU and regress. Citizenship of the rural transferring population is an important content of NEU [39], which also reflects the idea of human-centeredness of NEU. Citizenship of the rural transferring population is measured by 10 indicators, including the urbanization rate of resident population, urbanization rate of household registration, urban old-age insurance coverage rate, urban medical insurance coverage rate, the ratio of the number of people insured by unemployment insurance, the ratio of urban temporary residents, the ratio of general public budget expenditure on education, the ratio of general public budget expenditure on social security and employment, the ratio of general public budget expenditure on medical care, health care and family planning, and urban registered unemployment rate. The entropy weighting method is adopted to calculate the comprehensive score of citizenship of the rural transferring population, and E11 shows the coefficient of citizenship is 5.1312 on “quantity increase” of GI, and E12 shows that the coefficient of citizenship is 2.4748 on “quality improvement” of GI. Both of the regression coefficients pass the significance test, demonstrating that the benchmark regression outcomes are resilient.

#### v) Replacing dependent variable

Inaccurate measurement of the dependent variable may result in biased regression results. Although this paper uses applied green patents to measure GI, there may still be speculative behaviors or “policy dividend” behaviors that lead to the failure of the indicator to accurately measure GI. Therefore, this study changes the measurement of dependent variables, uses the total number of green patents granted per 10,000 people (NGIN) to measure the “quantity increase” of GI, and uses the total number of green invention patents granted per 10,000 people (NGNS) to measure the “quality improvement” of GI. The regression results are shown in Table 6. E13 shows that the regression coefficient of NEU on NGIN is 19.7964, and E14 shows that the regression coefficient of NEU on NGNS is 4.3227, both of which pass the test, demonstrating that the “quantity effect” and “quality effect” of NEU on GI are robust.

**Table 6.**
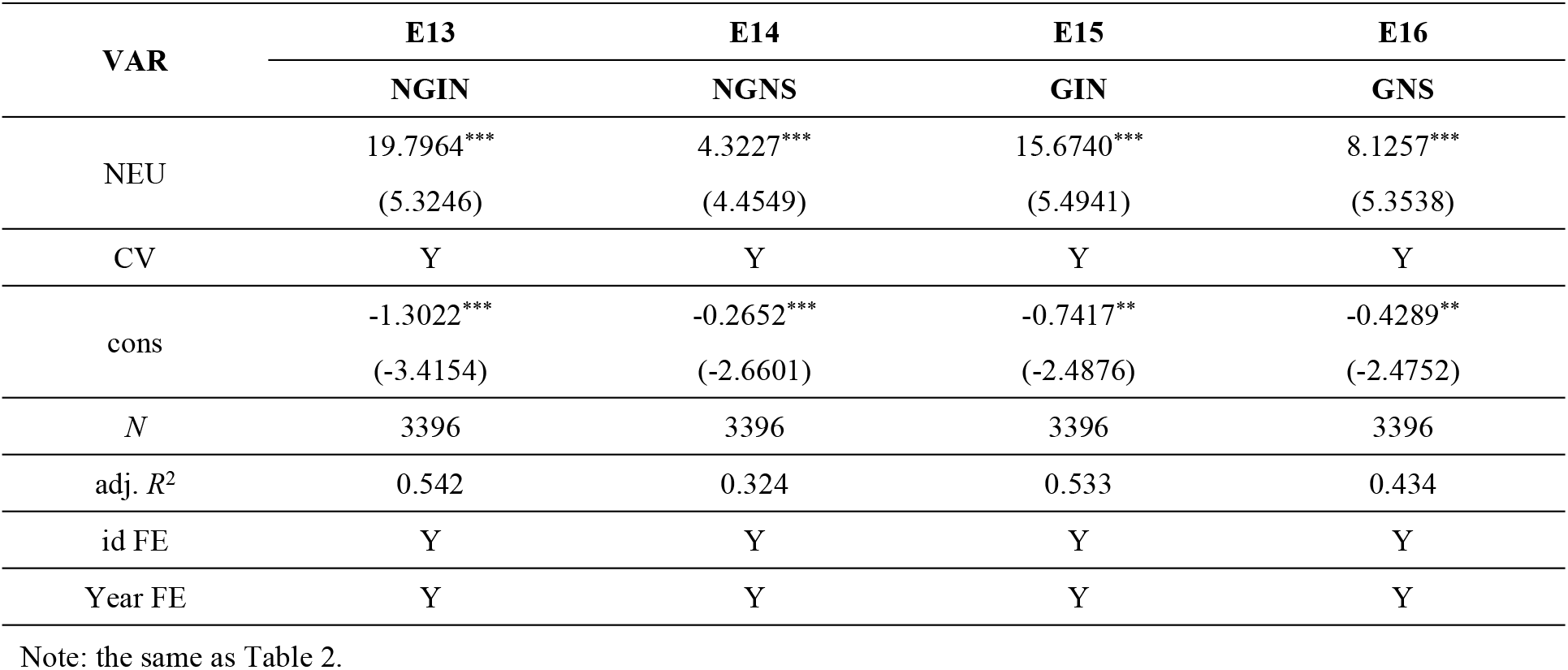
Results of Robustness Test III.

#### vi) Tailoring of the dependent variable

GIN and GNS are single indicators and may have extreme values. To mitigate the impact of extreme values on the regression results, the variables can be tailed [40]. E15 shows the coefficient of NEU is 15.6740, and E16 shows the coefficient of NEU is 8.1257; the regression coefficients are unchanged, which confirms the reliability of the benchmark regression.

### 4.3. Endogenous analysis

The above analysis indicates that NEU has a “quantity increase” and “quality improvement “effect on GI. However, in reality, cities with higher levels of GI tend to perform better in urbanization [41]. Therefore, there may be a two-way causality problem. In addition, there may be the problem of important variables closely related to NEU and GI being omitted in the model estimation, which may also lead to biased estimated coefficients. Therefore, this paper employs the instrumental variable methodology to examine endogeneity.

#### i) The independent variable lagged one period (LAG_NEU) is used as an instrumental variable

Considering that there is no reverse causality of GI on NEU in the previous period, the endogeneity of the regression is verified using LAG_NEU as an instrumental variable. The results are shown in Table 7. E17 shows that the coefficient of LAG_NEU on NEU is 0.4634, which meets the significance test, signifying that NEU lagged by one period has a positive correlation on NEU, which satisfies the condition of correlation. E18 shows that the coefficient of the estimation of NEU on GIN is 41.6357, and E19 shows that the coefficient is 23.3709, which meets the significance and Anderson canon. Corr. LM statistic is 603.319, p10, which passes the test for weak instrumental variables. The empirical results show that the conclusion remains robust after considering the endogeneity problem.

**Table 7.**
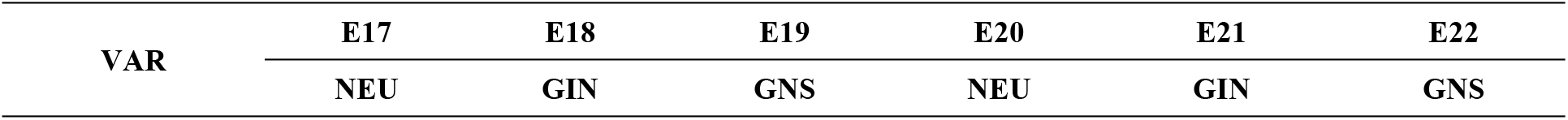
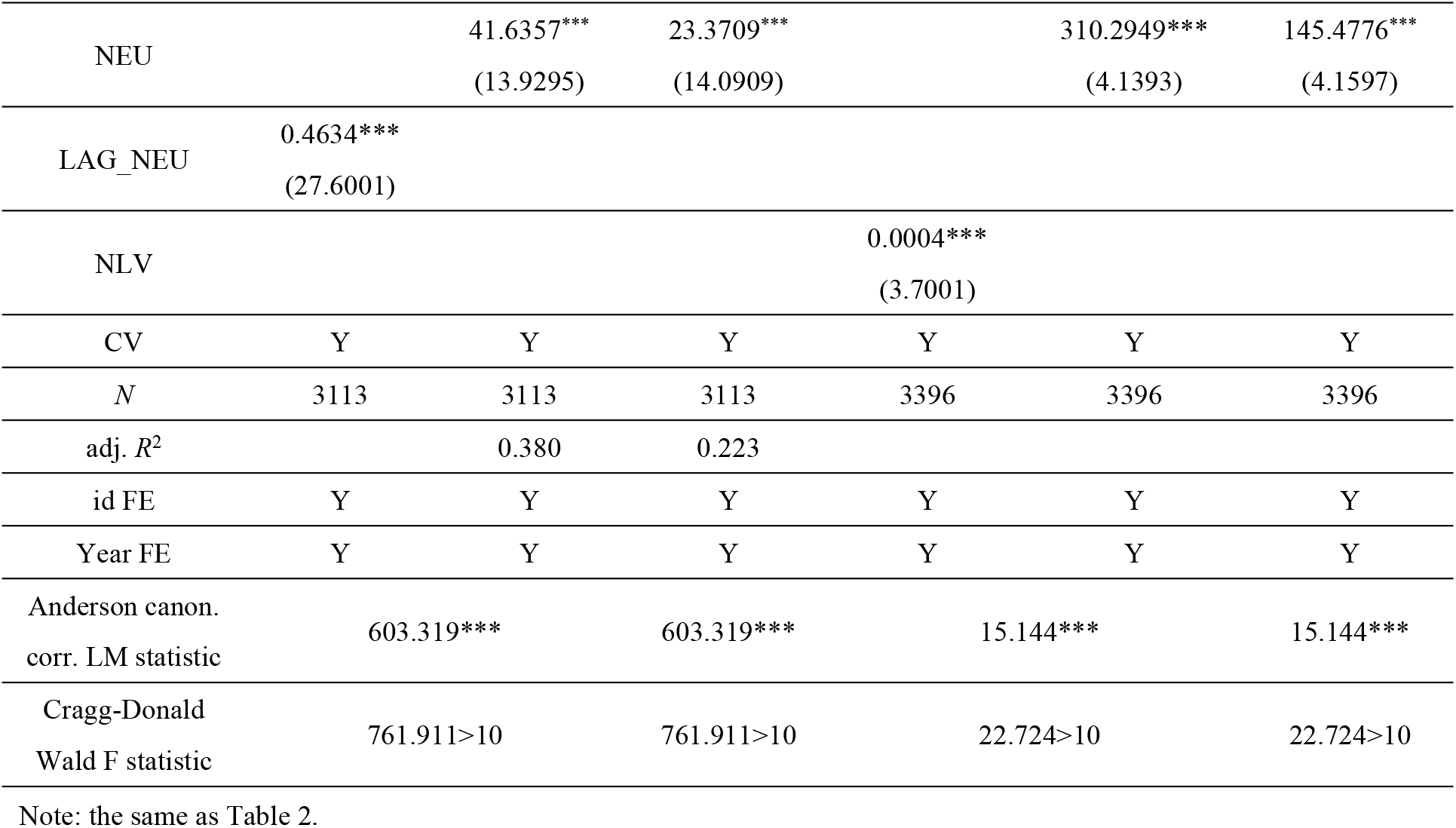
Results of endogeneity test.

#### ii) Night lighting value (NLV) as an instrumental variable

Night lighting value can reflect the human activities in a city; the higher the night lighting value, the higher the population concentration in the area [42], which can reflect the higher urbanization level in the area. Therefore, it can be assumed that there is a correlation between urban nighttime lighting values and NEU. Urban nighttime lighting has natural attributes, is generally directly observed by satellite remote sensing, is not affected by local government policy or economic indicators, and does not have a direct impact on GI, so NLV is an exogenous variable. Therefore, endogeneity is tested using NLV as an instrumental variable for NEU. E20 shows a regression coefficient of NLV on NEU is 0.0004; nighttime lighting values have a strong correlation with NEU. E21 shows a coefficient of NEU on GIN of 310.2949, and E22 shows a coefficient of NEU on GNS of 145.4776, both of which pass the test of significance. Anderson canon. Corr. LM statistic is 15.144, p<0.01, underidentification test passes. The Cragg-Donald Wald F=22.724>10 weak instrumental variable test passes. The empirical results indicate that the conclusions of the “quantity increase” and “quality improvement” effect of NEU on GI are valid after considering the endogeneity problem.

### 4.4. Moderating effects

NEU has both a “quantity increase” and a “quality improvement” effect on GI. However, it also varies greatly from city to city. This is closely related to the cities’ development status and positioning, as well as DIF, government support for technology, informatization, regional economic development and energy consumption. DIF has a lower service threshold compared to traditional finance, has a wider coverage of small-sized enterprises and low-income groups, improves the efficiency of financial services, and creates convenient conditions for the economic development of urban areas. Therefore, this study uses DIF as the main moderating variable and government technology support, informatization, regional economic development and energy consumption as the grouping variables to analyze the moderating effect of NEU on GI.

#### i) Moderating effect of DIF

DIF and the interaction term between NEU and DIF (cNEU×cDIF) are included in the regression model; the results are presented in Table 8. E23 shows that the coefficient of cNEU×cDIF is 0.0721, which passes the test of significance, and E24 shows that the coefficient of cNEU×cDIF is 0.0403, which indicates that DIF has a positive moderating effect. DIF has enhanced the “quantity increase” effect of NEU on GI and the “quality improvement” effect of NEU on GI. DIF reduces the financing threshold, attracts more funds to the field of GI, and meets the financial needs of GI technology research and development [43]. It can be seen that DIF can effectively alleviate the financing constraints faced by urbanization and GI, reduce the information friction of matching innovation factors, and enhance the marginal effect of urbanization on GI.

**Table 8.**
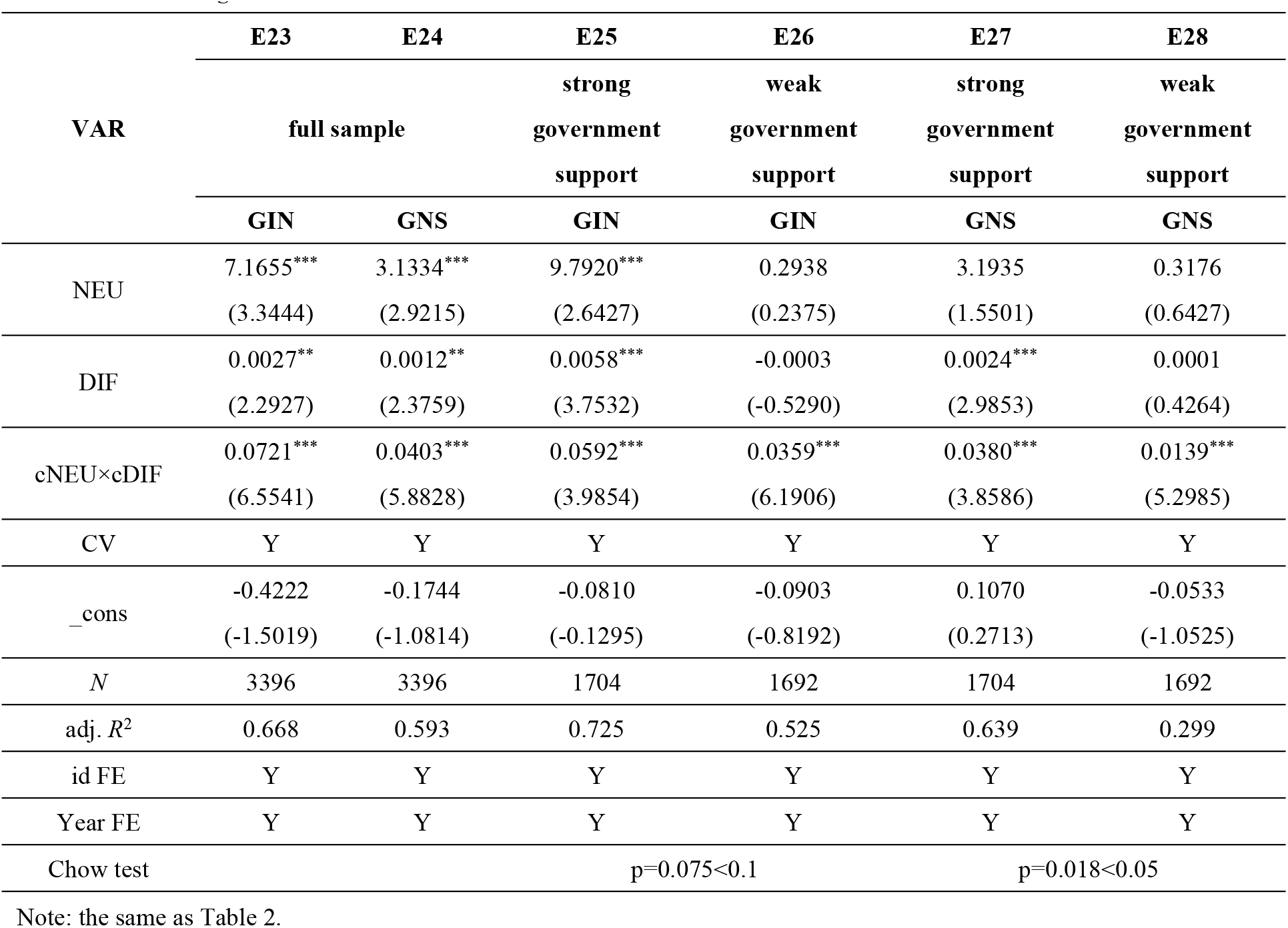
Moderating Effects Results I.

#### ii) Moderating effect of DIF and government technology support

Cities are grouped according to the median of GTS, and cities with GTS greater than the median are “strong government support cities”, while cities with GTS less than the median are “weak government support cities “. E25 shows that the regression coefficient of cNEU×cDIF is 0.0592, and E26 shows that the regression coefficient of cNEU×cDIF is 0.0359, which both meet the importance criterion. The difference between group coefficients of the interaction term is tested, p=0.075<0.1. This indicates that the moderating influence of DIF is more pronounced in strong government-support cities and weaker in weak government-support cities. DIF can better strengthen the “quantity increase” effect of NEU on GI in strong government-supported cities. E27 shows the coefficient of cNEU×cDIF is 0.0380, and E28 shows the coefficient of cNEU×cDIF is 0.0139, which both meet the significance criterion. Difference in the coefficients between groups of interaction term coefficients is tested, p=0.018<0.05. This indicates that the moderating effect of DIF is stronger in strong government-support cities, while the moderating effect of DIF in weak government-support cities is relatively weaker. This is because in strong government support cities, industries are better integrated with the digital economy [44], and DIF can better strengthen the “quality improvement” effect of NEU on GI. Government support optimizes the resource allocation capacity of digital finance, decreases the incremental expense of financial services, lowers the risk of enterprise R&D and innovation [45], and helps accelerate the diffusion of green technologies in urbanization, which improves the performance of GI.

#### iii) Moderating effects of DIF and informatization

Cities are grouped according to the median of INT, and cities with INT greater than the median are “high informatization cities”, while cities with INT less than the median are “low informatization cities”. The results are presented in Table 9. E29 shows the coefficient of cNEU×cDIF is 0.0596, and E30 shows the coefficient of cNEU×cDIF is 0.0355, both of which pass the significance test. The difference in interaction terms between groups is tested, p=0.008<0.01. In high-informatization cities, the moderating effects of DIF are stronger. In contrast, in low-informatization cities, the moderating effect of DIF is relatively weaker, indicating that DIF in high-informatization cities can better strengthen the “quantity increase” effect of NEU on GI. E31 shows the coefficient of cNEU×cDIF is 0.0426, and E32 shows the coefficient of cNEU×cDIF is 0.0193, both of which passed the significance test. The difference in interaction terms between groups is tested, p=0.036<0.05. This indicates that the moderating effect of DIF is stronger in high-informatization cities and that DIF can better strengthen the “quality improvement” effect of NEU on GI. This also verifies that DIF and informatization do not act independently but are mutually reinforcing processes. Digital finance relies on informatization infrastructure, and informatization needs financial support [46]; high informatization cities can more efficiently match digital financial resources with green technology needs, reduce transaction costs, capital, technology, and talent can flow more efficiently to the field of GI, which strengthening the connection between NEU and GI.

**Table 9.** Moderating Effects Results II.

| VAR | E29 | E30 | E31 | E32 |
| --- | --- | --- | --- | --- |
|  | high informatization | low informatization | high informatization | low informatization |
|  | cities | cities | cities | cities |
|  | GIN | GIN | GNS | GNS |
| NEU | 4.0175<br>(0.9121) | 4.0627***<br>(3.5958) | 0.0016<br>(0.0006) | 1.9512***<br>(3.2639) |
| DIF | 0.0026<br>(1.1749) | 0.0009**<br>(2.3993) | 0.0008<br>(0.7303) | 0.0006***<br>(2.6675) |
| $cNEU \times cDIF$ | 0.0596***<br>(3.5925) | 0.0355***<br>(4.9196) | 0.0426***<br>(3.3800) | 0.0193***<br>(5.2191) |
| CV | Y | Y | Y | Y |
| _cons | 0.1164<br>(0.1782) | -0.4047***<br>(-2.9016) | 0.4734<br>(1.0499) | -0.2417***<br>(-2.8112) |
| N | 1700 | 1696 | 1700 | 1696 |
| adj. $R^2$ | 0.690 | 0.503 | 0.629 | 0.343 |
| id FE | Y | Y | Y | Y |
| Year FE | Y | Y | Y | Y |
| Chow test | $p=0.008<0.01$ | | $p=0.036<0.05$ | |

Note: the same as Table 2.

#### iv) Moderating effects of DIF and regional economic development

Cities are grouped according to the median per capita GDP (AGDP), and cities with AGDP greater than the median are “developed cities”, while cities with AGDP less than the median are “underdeveloped cities”. The results are presented in Table 10. The moderating variables and interaction terms are added to the regression model for analysis. The coefficient of cNEU×cDIF is 0.0571 in the “developed cities” group. The interaction term is 0.0270 in the “underdeveloped cities” group, and the differences in the interaction term coefficients between the groups are tested, p=0.080<0.1. This indicates that the moderating effect of DIF is more pronounced in the “developed cities” group; DIF better enhances the “quantity increase” effect of NEU on GI. The moderating variables and cNEU×cDIF are added to the model of quality improvement effect. The coefficient of cNEU×cDIF is 0.0378 in the “developed cities” group and 0.0118 in the “underdeveloped cities” group; the differences of the interaction terms coefficients between the groups are tested, p=0.077<0.1. This suggests that the moderating effect of DIF is stronger in the “developed cities” group and that DIF better enhances the “quality improvement” effect of NEU on GI. This is because the synergistic effect of the “digital capital market” will be formed when DIF interacts with economic development. DIF accurately matches the demand for capital and accelerates the commercialization of green technologies [47]. At the same time, economic development brings about the market scale, enhances the absorption capacity of technology, and further amplifies the benefits of GI. This synergy significantly enhances the positive impact of NEU on GI.

**Table 10.**
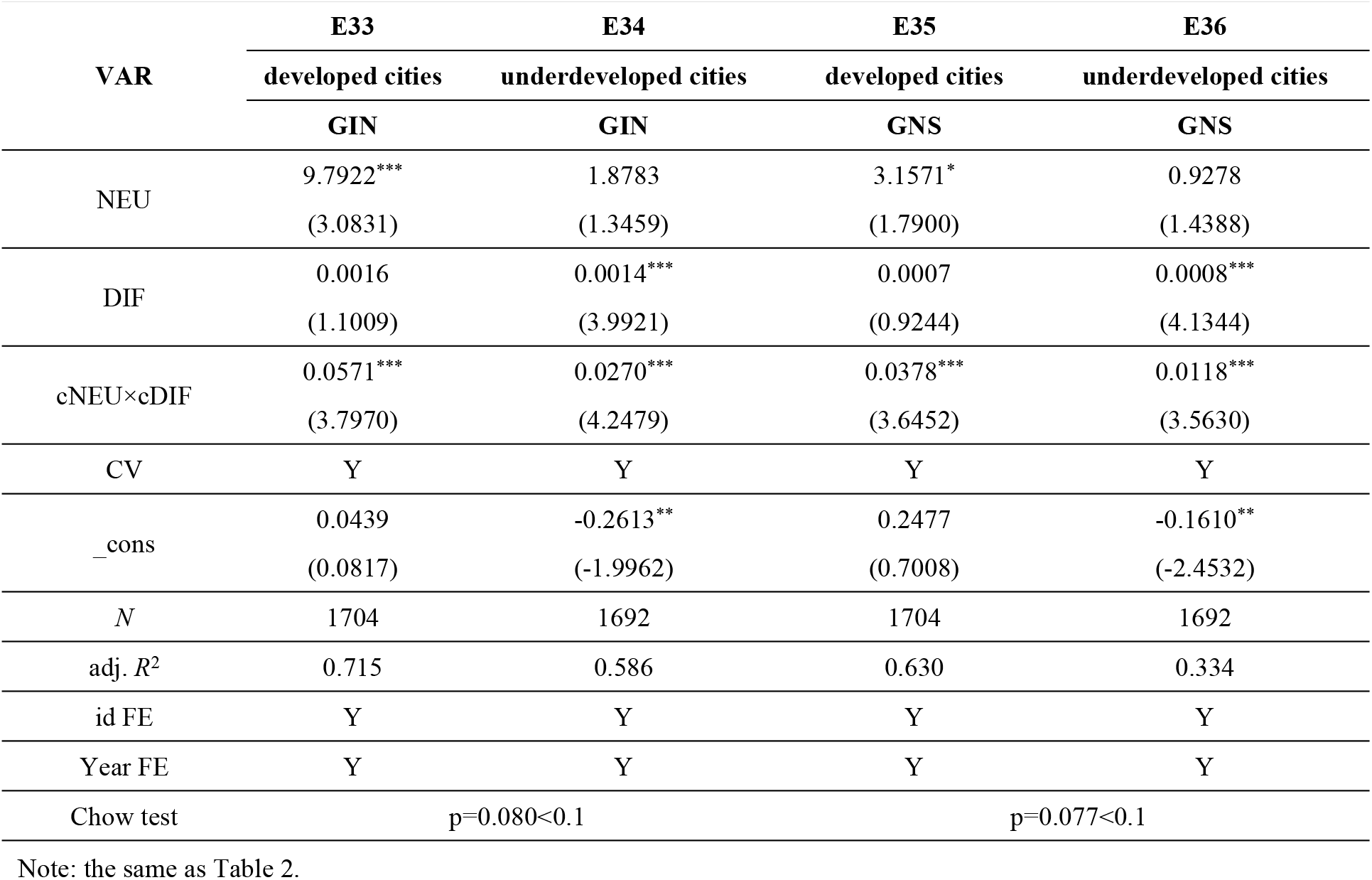
Moderating Effects Results III.

| VAR | E33 | E34 | E35 | E36 |
| --- | --- | --- | --- | --- |
|  | developed cities | underdeveloped cities | developed cities | underdeveloped cities |
|  | GIN | GIN | GNS | GNS |
| NEU | 9.7922***<br>(3.0831) | 1.8783<br>(1.3459) | 3.1571*<br>(1.7900) | 0.9278<br>(1.4388) |
| DIF | 0.0016<br>(1.1009) | 0.0014***<br>(3.9921) | 0.0007<br>(0.9244) | 0.0008***<br>(4.1344) |
| $cNEU \times cDIF$ | 0.0571***<br>(3.7970) | 0.0270***<br>(4.2479) | 0.0378***<br>(3.6452) | 0.0118***<br>(3.5630) |
| CV | Y | Y | Y | Y |
| _cons | 0.0439<br>(0.0817) | -0.2613**<br>(-1.9962) | 0.2477<br>(0.7008) | -0.1610**<br>(-2.4532) |
| N | 1704 | 1692 | 1704 | 1692 |
| adj. $R^2$ | 0.715 | 0.586 | 0.630 | 0.334 |
| id FE | Y | Y | Y | Y |
| Year FE | Y | Y | Y | Y |
| Chow test | $p=0.080<0.1$ | | $p=0.077<0.1$ | |

Note: the same as Table 2.

#### v) Moderating effects of DIF and energy consumption

Significant disparities exist in industrial institutions, resource endowments, technological levels, industrial characteristics, energy policies, and economic development models in different cities, which inevitably lead to differences in energy consumption and energy utilization efficiency. Energy consumption will also have an impact on GI in each city. Cities are grouped according to the median energy consumption (ENC). Cities with ENC greater than the median are “high-energy-consuming cities”, and those with ENC less than the median are “low-energy-consuming cities”. The coefficient of cNEU×cDIF is 0.0644 in “high-energy-consuming cities” and 0.0438 in the “low-energy-consuming cities” (table 11), and the test of difference of the coefficients between the groups shows that the difference is significant. This suggests that the moderating effect of DIF is stronger in “high-energy-consuming cities” and that DIF better enhances the “quantity increase” effect of NEU on GI. The moderating effect of NEU on quality improvement of GI shows that the coefficient of cNEU×cDIF is 0.0390 in “high-energy-consuming cities” and 0.0211 in “low-energy-consuming cities”; the difference between the groups is significant. This suggests that the moderating effect of DIF is stronger in “high-energy-consuming cities”, and DIF better strengthens the “quality improvement” effect of NEU on GI. Because the development of DIF provides financial support for GI, and the increase in energy consumption stimulates the demand for GI [48], a positive feedback effect occurs when DIF interacts with energy consumption.

**Table 11.**
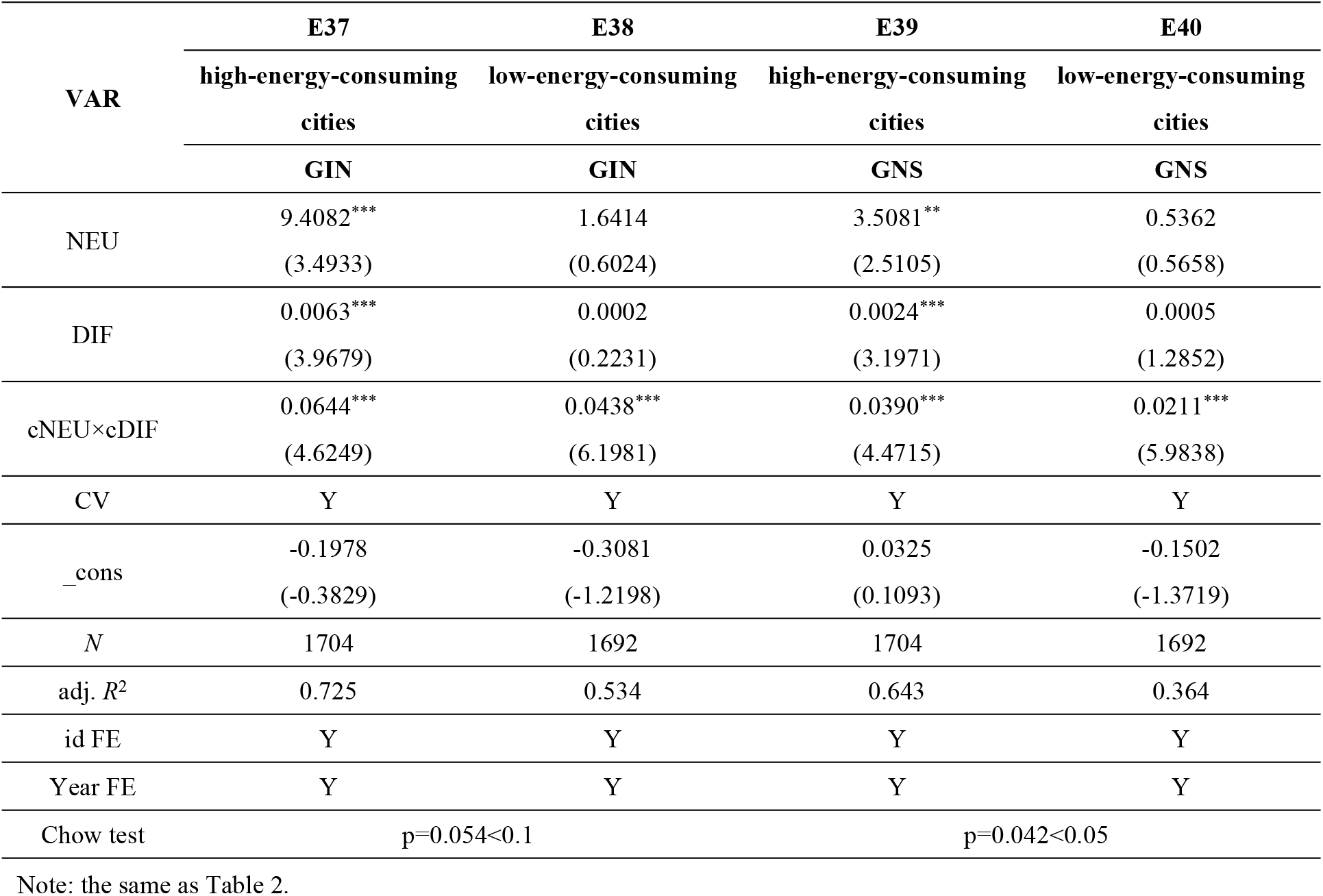
Moderating Effects Results IV.

| VAR | E37 | E38 | E39 | E40 |
| --- | --- | --- | --- | --- |
|  | high-energy-consuming | low-energy-consuming | high-energy-consuming | low-energy-consuming |
|  | cities | cities | cities | cities |
|  | GIN | GIN | GNS | GNS |
| NEU | 9.4082***<br>(3.4933) | 1.6414<br>(0.6024) | 3.5081**<br>(2.5105) | 0.5362<br>(0.5658) |
| DIF | 0.0063***<br>(3.9679) | 0.0002<br>(0.2231) | 0.0024***<br>(3.1971) | 0.0005<br>(1.2852) |
| cNEU×cDIF | 0.0644***<br>(4.6249) | 0.0438***<br>(6.1981) | 0.0390***<br>(4.4715) | 0.0211***<br>(5.9838) |
| CV | Y | Y | Y | Y |
| _cons | -0.1978<br>(-0.3829) | -0.3081<br>(-1.2198) | 0.0325<br>(0.1093) | -0.1502<br>(-1.3719) |
| N | 1704 | 1692 | 1704 | 1692 |
| adj. R<sup>2</sup> | 0.725 | 0.534 | 0.643 | 0.364 |
| id FE | Y | Y | Y | Y |
| Year FE | Y | Y | Y | Y |
| Chow test | p=0.054<0.1 |  | p=0.042<0.05 |  |

Note: the same as Table 2.

## 5 Further analysis

To examine whether the impact of NEU on GI exhibits a nonlinear relationship, a panel threshold effect model is constructed:

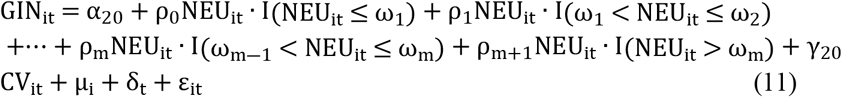

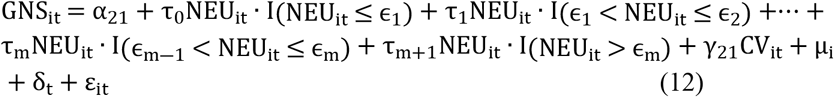

ρ is the coefficient of NEU, I(·)is an indicator function, ω and ϵ are threshold values.

The threshold variable is government green support (GES), which is measured by the ratio of fiscal environmental protection expenditure to fiscal general budget expenditure. First, we need to determine the number of thresholds. This paper uses Bootstrap to test, and the estimation results are presented in Table 12. When GIN is the dependent variable, the single-threshold value and the double-threshold value are significant, and the triple-threshold is not significant. It indicates that there is a double-threshold effect of government green support. When GNS is the dependent variable, both the single-threshold value and double-threshold value are significant, and the triple-threshold is not significant, indicating that there is also the double-threshold effect of government green support. The threshold values of government green support are 0.0081 and 0.0074.

**Table 12.** Threshold Test Results.

| Threshold | dependent variable: GIN |  |  |  | dependent variable: GNS |  |  |  |
| --- | --- | --- | --- | --- | --- | --- | --- | --- |
|  | RSS | MSE | Fstat | Prob | RSS | MSE | Fstat | Prob |
| Single | 1016.6010 | 0.3004 | 16.72 | 0.0133 | 310.2835 | 0.0917 | 23.40 | 0.0000 |
| Double | 1012.2516 | 0.2991 | 14.54 | 0.0100 | 308.4852 | 0.0912 | 19.73 | 0.0067 |
| Triple | 1010.6460 | 0.2987 | 5.38 | 0.6000 | 308.0675 | 0.0910 | 4.59 | 0.6633 |

We further analyze the non-linear effects of NEUs on the “quantity increase” and “quality improvement” of GI under government green support. The estimation results are presented in Table 13. When the government green support is less than 0.0074, the coefficient of NEU on the “quantity increase” of GI is 21.5998, which is significant. When 0.0074<GES≤0.0081, the coefficient of NEU on the “quantity increase” of GI is 22.7229, which is significant. When GES>0.0081, the coefficient of NEU on the “quantity increase” of GI is 23.1905, which is significant. This means that as the government investment in environmental protection increases, it will be conducive for NEU to further exert the “quantity increase” effect on GI, and the government green support makes the “quantity increase” effect show the trend of increasing marginal efficiency. The government’s regulation of environmental protection, limitations on high-emission and energy-intensive sectors, and other initiatives facilitate the development of innovative business models, including pollution monitoring devices and solid waste resource use [49]. The government’s augmented investment in environmental protection will facilitate the swift advancement of green and eco-friendly industries. NEU further stimulates the market application of environmental protection technology and innovation power. The rapid development of the green environmental protection industry also creates important conditions for talent gathering and urbanization transformation. Therefore, as the government’s green support increases, it is conducive to expanding the quantity increase effect of NEU on GI.

**Table 13.** Threshold Effect Estimation Results.

| VAR | E41 | E42 |
| --- | --- | --- |
|  | GIN | GNS |
| CV | Y | Y |
| NEU($GES \leq 0.0074$) | 21.5998***<br>(5.8090) | 10.9981***<br>(5.2460) |
| NEU($0.0074 < GES \leq 0.0081$) | 22.7229***<br>(6.0566) | 11.7606***<br>(5.3457) |
| NEU($GES > 0.0081$) | 23.1905***<br>(6.2748) | 12.0378***<br>(5.4309) |
| _cons | -1.2133***<br>(-3.0678) | -0.6254***<br>(-2.7349) |
| N | 3396 | 3396 |
| adj. $R^2$ | 0.523 | 0.417 |
| id FE | Y | Y |
| Year FE | Y | Y |

Note: the same as Table 2.

Similarly, When GES≤0.0074, the coefficient of NEU on “quality improvement” of GI is 10.9981, which is significant. When 0.0074<GES≤0.0081, the coefficient of NEU on the “quality improvement” of GI is 11.7606, which is significant. When GES>0.0081, the coefficient of NEU on the “quality improvement” of GI is 12.0378, which is significant; as the government’s investment in environmental protection increases, it forces the traditional manufacturing industries to transform its manufacturing processes, guiding it towards enhanced efficiency and sustainable transformation [50], realizing the technological transformation of high-energy-consuming industries. Simultaneously, the advancement of solid waste resource utilization technology in urbanization construction and the enhancement of resource recycling efficiency would contribute to improving the quality improvement of GI.

## 6. Discussion

### 6.1. Summary

During the Fourteenth Five-Year Plan period, China is in a period of rapid urbanization, and urbanization development faces both opportunities and challenges. Against this background, China put forward the NEU strategy to coordinate resources and elements in order to transform the urban development mode, promote the growth and concentration of innovative talents, enhance the urban innovation capacity and innovation effectiveness, and help the sustainable development of urban and regional economies. This paper empirically analyzes the “quantity increase” and “quality improvement” effects of NEU on GI by adopting urban panel data from 2011 to 2022. It was found that NEU not only has a positive contribution to the quantity increase of GI but also plays a positive role in the quality improvement of GI. The robustness test and the endogeneity test verify that the conclusion is valid. DIF further amplifies the dividends of NEU and plays a promoting effect on GI. The interaction of DIF and government support, informatization, regional economic development, and energy consumption also amplify the dividends of NEU and exert strong “quantity increase” and “quality improvement” effects on GI. The impact of NEU on GI has a nonlinear relationship, and the stronger the government’s green support is, the more conducive it is for NEU to exert “quantity increase” and “quality improvement” effects on GI.

### 6.2. Recommendations

(1) This study finds that NEU not only has a positive role in promoting quantity increase of GI but also plays a positive role in quality improvement of GI. Therefore, we need to scientifically plan the functional zoning of the cities, rationally plan the distribution and layout of green industries and high energy-consuming industries, plan specialized GI industry clustering zones, and create GI spatial carriers so as to create favorable conditions for GI. Governments should guide the transfer of population into green industries and ensure the talent demand of green industries, build efficient urban form, improve urban land utilization efficiency, reduce urban congestion and energy consumption, increase the construction of urban infrastructure, accelerate the adjustment and optimization of the urban energy structure, increase investment in green technology, guide enterprises to carry out green product innovation, guide the transferring population to engage in GI and recognize and reward enterprises, research institutions and individuals that have made outstanding contributions to green employment and GI. (2) This study finds that DIF further amplifies the dividends of NEU and exerts a stronger GI effect. Therefore, we need to build a green finance digital service platform to provide accurate decision support for financial institutions, clarify the demand for GI, develop an innovative DIF system, enrich green credit products, promote green financial product innovation, and meet the diversified needs of enterprises and research institutions. Government departments can establish a green financial information-sharing platform, encourage the cross-regional flow of green financial resources, promote the transfer of funds from financially developed regions to less developed regions, and realize cross-regional docking of funds, resources and markets. The management department should fully leverage the synergistic effects of DIF and government technological support, informatization, regional economic development, and energy efficiency to unleash the dividends of NEU further and enhance its service efficiency. (3) Government departments should ensure the proportion of annual financial expenditures for environmental protection, provide loan subsidies to GI enterprises or projects to reduce their financing costs, set up a GI guidance fund, actively introduce social capital to participate in NEU and GI and attract enterprises to participate in the construction of urban eco-parks, eco-city construction. Social capital entities participating in green innovation should be included in the environmental credit evaluation system and given appropriate preferential treatment in government procurement and project bidding.

## Supporting information

Data(XLS)

## Author Contributions

Conceptualization, L.F. and C.L.; methodology, L.F. and C.L.; validation,L.F. and C.J.; formal analysis, L.F. and C.J.; investigation, L.F. and C.L.; resources, L.F.;writing—original draft preparation, L.F. and C.L.; visualization, L.F. and C.J.; supervision, L.F.and C.L.; project administration, L.F. and C.J. All authors have read and agreed to the published version of the manuscript.

